# Exploiting the unwanted: sulphate reduction enables phosphate recovery from energy-rich sludge during anaerobic digestion

**DOI:** 10.1101/584904

**Authors:** Celine Lippens, Jo De Vrieze

**Affiliations:** Center for Microbial Ecology and Technology (CMET), Ghent University, Coupure Links 653, B-9000 Gent, Belgium

**Author notes:** Correspondence to: Jo De Vrieze, Ghent University; Faculty of Bioscience Engineering; Center for Microbial Ecology and Technology (CMET); Coupure Links 653; B-9000 Gent, Belgium; Webpage: www.cmet.ugent.be.

**Keywords:** Biogas, methanogenesis, resource recovery, sulphate reducing bacteria

## Abstract

Anaerobic digestion is shifting from a single-purpose technology for renewable energy recovery from organic waste streams to a process for integrated resource recovery. The valorisation of high-rate energy- and phosphorus-rich sludge creates the opportunity for their combined recovery. This phosphate is present in a precipitated form in the sludge, and its release into the liquid phase is an important issue before recovery can be achieved. The objective of this research was to exploit the “unwanted” sulphate reduction process for the release of phosphate into the liquid phase during anaerobic digestion, thus, making it available for recovery. Two different treatments were considered, *i.e*., a control digester and a digester to which sulphate was added, each operated in triplicate for a period of 119 days. The control digester showed stable methane production at 628 ± 103 mL CH_4_ L^−1^ d^−1^, with a feedstock COD (chemical oxygen demand) conversion efficiency of 89.5 ± 14.6 %. In contrast, the digester with sulphate addition showed a 29.9 ± 15.3 % decrease in methane production, reaching an “inhibited steady state”, but phosphate release into the liquid phase increased with a factor 4.5, compared to the control digester. This inhibited steady state coincided with a clear shift from a Methanosaetaceae to a Methanosarcinaceae dominated methanogenic community. Overall, the sulphate reduction process allows phosphate release during the anaerobic digestion process, yet, at the cost of a reduced methane production rate.

## 1. Introduction

Anaerobic digestion (AD) has been a key technology for the recovery of renewable energy from organic waste streams for decades. Initially, however, the main purpose of AD was the stabilisation of organic waste streams to avoid environmental pollution (Acosta and De Vrieze 2018). The ability to use the energy-rich methane in a combined heat and power (CHP) unit for electricity and heat production quickly allowed the transition of AD from a waste treatment technology to an integrated system for renewable energy recovery. Different organic waste streams, such as animal manure (Holm-Nielsen et al. 2009), waste activated sludge (Appels et al. 2008), the organic fraction of municipal solid waste (Hartmann and Ahring 2006) have been valorised through AD, either as such or through co-digestion with other waste streams (Björn et al. 2017, Mata-Alvarez et al. 2011). In the framework of the current transition from “waste-to-energy” to “waste-to-resource”, the recovery of nutrients, in addition to energy, has become more and more pressing to (1) safeguard natural resources and (2) ensure long-term economic viability of the AD process.

The recovery of nutrients in refined products through AD has been demonstrated through numerous technologies in multiple configurations. Stripping/absorption is a well-established method for ammonia recovery in AD, either as pre-treatment (Bonmatí and Flotats 2003, Zhang et al. 2012), in side stream (Pedizzi et al. 2017, Serna-Maza et al. 2014), or post-treatment technology (Bonmatí and Flotats 2003, Gustin and Marinsek-Logar 2011). An alternative approach involves the electrochemical recovery of ammonium and other cations *via* a cation exchange membrane (Desloover et al. 2012), thus, allowing their recovery in a “clean” stream (De Vrieze et al. 2018). Both technologies are often combined for efficient ammonia recovery, as electrochemical extraction, followed by stripping/absorption enables the creation of a continuous concentration gradient (Desloover et al. 2015, Zhang and Angelidaki 2015).

The recovery of phosphorus in combination with AD is often problematic, because phosphates precipitate with multivalent cations, such as Ca^2+^, Mg^2+^ and Fe^2+^, say in the case of waste activated sludge (De Vrieze et al. 2016). Hence, the release and recovery of phosphorus from waste activated sludge requires alternative approaches, such as a microwave treatment (Liao et al. 2005), a free ammonia-based pre-treatment (Xu et al. 2018), or pressurized AD (Latif et al. 2018). This allows the release of phosphate into the liquid phase, and the potential for subsequent recovery either through (1) struvite precipitation, to be used as slow-release fertilizer (Li et al. 2019, Vaneeckhaute et al. 2018), (2) electrodialysis, using an anion exchange (Ebbers et al. 2015) or bipolar (Shi et al. 2018) membrane system, or (3) a combination thereof (Zhang et al. 2013). These different technologies for the release of phosphate from the solid phase, however, require the input of chemicals and/or a coincide with an additional energy cost per unit of phosphorus released. Given the low and variable global market value of phosphorus, *i.e*., € 350-1200 tonne^−1^ P for phosphate rock with a P_2_O_5_ content of 30% since 2010, alternative low-cost strategies for phosphate release should be targeted (Mayer et al. 2016).

Combining AD of waste activated sludge with *in situ* sulphate reduction by sulphate reducing bacteria could be an alternative approach for phosphate release into the liquid phase with no additional requirements in terms of energy or chemicals by using sulphate-rich waste streams as co-feedstock, such as vinasse or paper mill wastewater (Pokhrel and Viraraghavan 2004, Rodrigues Reis and Hu 2017). As the solubility product of multivalent cations with sulphides is conventionally lower than with phosphates, the *in situ* formation of sulphides could release phosphate into the liquid phase. The reduction of sulphate to sulphide during AD could, however, negatively impact methane production due to (1) competition between sulphate reducing bacteria and methanogens for “reducing power”, (2) direct inhibition of methanogens, due to H_2_S toxicity, and (3) reduced trace metal bioavailability, due to precipitation with sulphides (Karhadkar et al. 1987, Paulo et al. 2015). Hence, accurate control of this “unwanted” sulphate reduction process, by monitoring the ingoing sulphate concentration and H_2_S content in the biogas, is essential to achieve a long-term stable integrated process of methane production and phosphorus release.

The key objective of this study was to obtain integrated energy recovery, through the production of biogas, and phosphate release from high-rate P-rich activated sludge during AD without the need for additional chemicals. Sulphate reduction by sulphate reducing bacteria, which is commonly considered an “unwanted process”, was carefully steered during AD operation to maximise the release of phosphate into the liquid phase, whilst limiting the impact of the sulphate reduction process on methane production.

## 2. Material and methods

### 2.1. Inoculum and feedstock

The high-rate activated sludge (A-sludge) that was used as feedstock during operation of the digesters was obtained as a single batch from the A-stage of the wastewater treatment plant Nieuwveer, Breda, the Netherlands (Table 1). The A-sludge was stored at 4°C until use. This wastewater treatment plant was operated at a short sludge retention time (< 2 days) to maximise the recovery of organics, according to the Adsorptions-Belebungsverfahren or AB-system principles (Boehnke et al. 1997, Meerburg et al. 2016). The inoculum for the anaerobic digesters was obtained from the sludge digesters at the full-scale wastewater treatment plant the Ossemeersen, Ghent, Belgium (Table 2).

**Table 1.**
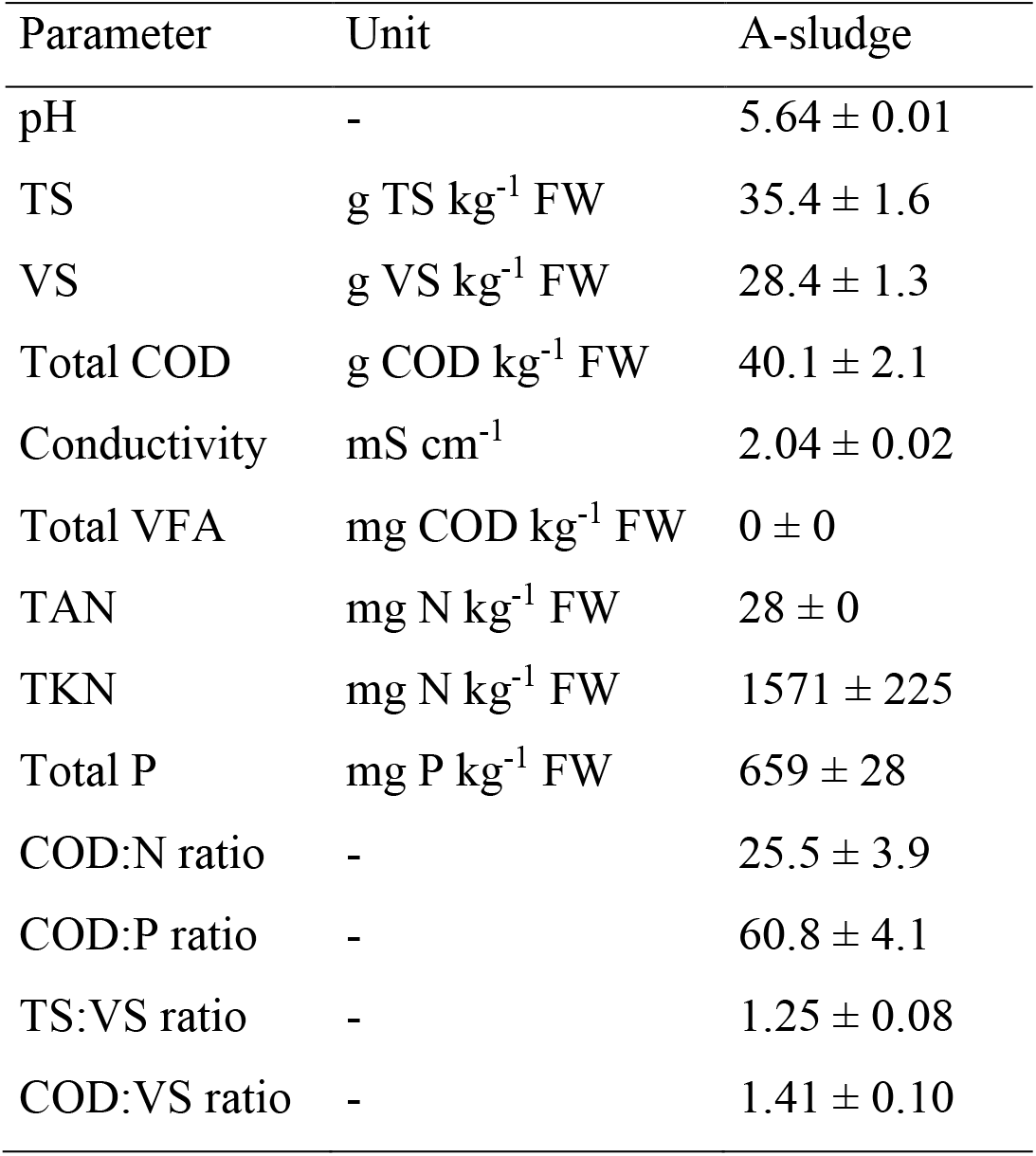
Main characteristics of the high-rate energy-rich A-sludge (n=3). TS = total solids, VS = volatile solids, COD = chemical oxygen demand, VFA = volatile fatty acids, TAN = total ammonia nitrogen, TKN = Kjeldahl nitrogen, FW = fresh weight.

**Table 2.**
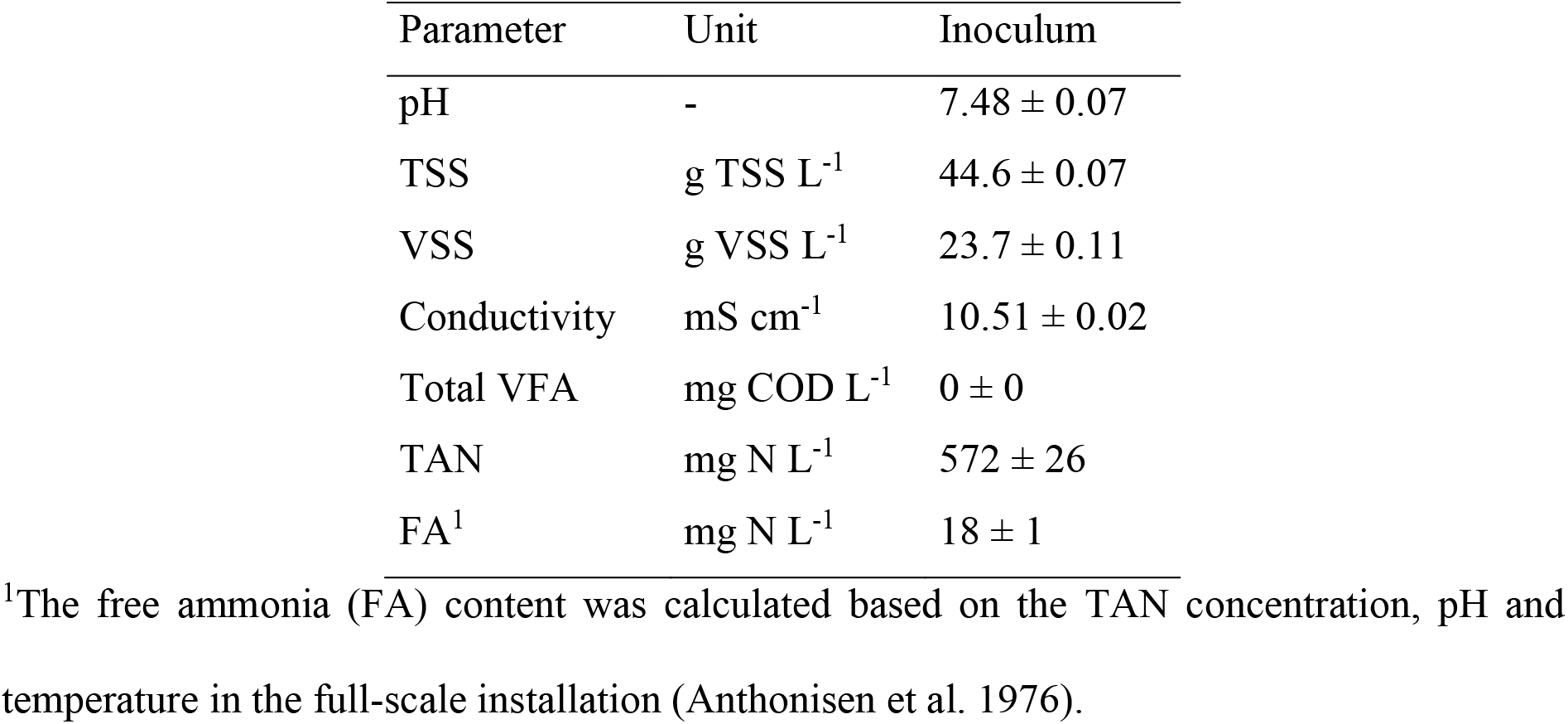
Main characteristics of the inoculum sludge (n=3). TSS = total suspended solids, VSS = volatile suspended solids, COD = chemical oxygen demand, TAN = total ammonia nitrogen, VFA = volatile fatty acids, FA = free ammonia nitrogen.

### 2.2. Experimental design and operation

Six Schott bottles with a total volume of 1 L and a working volume of 800 mL were operated as lab-scale anaerobic digesters. The bottles were sealed with air-tight rubber stoppers and connected to a water displacement system *via* gas-tight PVC tubing to monitor biogas production (Figure S1). The liquid in this system was kept at a pH < 4.3 to avoid CO_2_ in the biogas from dissolving. A Laboport^®^ vacuum pump (KNF Group International, Aartselaar, Belgium) and glass sampling tube of 250 mL (Glasgerätebau Ochs, Lenglern, Germany) were used to collect samples for biogas composition analysis. The initial inoculum biomass concentration in each digester was fixed at 10 g VSS L^−1^ (volatile suspended solids) by diluting the inoculum with tap water. The digesters were operated in a continuous stirred tank reactor mode with manual mixing, thus, the solids and hydraulic retention times were identical. Mesophilic conditions were maintained by operating the digesters in a temperature-controlled room at 34 ± 1°C. Digestate removal and feeding was carried out manually in fed-batch mode three times per week.

A start-up period of 14 days was implemented during which a sludge retention time of 40 days and an average organic loading rate of 1 g COD L^−1^ d^−1^ (chemical oxygen demand) were applied to allow adaptation of the microbial community in the inoculum to the new feedstock. From day 15 till day 119 (end of the experiment), a hydraulic retention time of 20 days and an average organic loading rate of 2 g COD L^−1^ d^−1^ were used. Until day 41, all six reactors were operated under identical conditions. From day 42 on, three biological replicates were subjected to sulphate addition at a predefined fixed sulphur to phosphorus molar ratio S:P of 2 by adding Na_2_SO_4_ to the feed (Sulphate digester). The other three biological reactors served as control digester with no sulphate addition (Control digester).

Biogas production and composition were monitored three times per week, together with the digester pH. Biogas production values were reported at standard temperature (273 K) and pressure (101325 Pa) conditions (STP). The sulphate, phosphate, sodium, total ammonium and volatile fatty acids (VFA) concentrations were measured on a weekly basis. The free ammonia (NH3) concentration was calculated based on the pH and total ammonia concentration (Anthonisen et al. 1976). The overall salinity in the digesters was estimated through a weekly measurement of the conductivity. Samples for microbial community analysis were taken on day 0 (inoculum and A-sludge), and day 42, 82 and 119 from each digester, and stored at −20°C until DNA extraction was performed.

### 2.3. Microbial community analysis

The frozen samples were subjected directly to DNA extraction with the ZymoBIOMICS™ DNA Miniprep Kit (Zymo Research, Irvine, CA, USA), using a PowerLyzer^®^ 24 Bench Top Bead-Based Homogenizer (MO BIO Laboratories, Inc, Carlsbad, CA, USA), and following the manufacturer’s instructions. The quality of the DNA extracts was validated with agarose gel electrophoresis and through PCR analysis using the universal bacterial primers 341F (5’-CCTACGGGNGGCWGCAG) and 785Rmod (5’-GACTACHVGGGTATCTAAKCC) that target the V3-V4 region of the 16S rRNA gene (Klindworth et al. 2013), following the protocol of Boon et al. (2002). The samples were sent to BaseClear B.V., Leiden, The Netherlands, for Illumina amplicon sequencing of the V3-V4 region of the 16S rRNA gene of the bacterial community on the MiSeq platform with V3 chemistry. The amplicon sequencing and data processing are described in detail in SI (S2). Real-time PCR analysis was carried out to quantify total bacteria, the methanogenic orders Methanobacteriales and Methanomicrobiales, and the methanogenic families Methanosaetaceae and Methanosarcinaceae, as described in SI (S3).

### 2.4. Statistical analysis

A table with the relative abundances of the different bacterial OTUs (operational taxonomic units), together with their taxonomic assignment (Supplementary file 2) was generated following the amplicon data processing. All statistical analysis were carried out in R Studio version 3.3.1 (http://www.r-project.org) (R Development Core Team 2013). First, a repeated measures analysis of variance (ANOVA, *aov* function) was used to validate that the biological replicates showed no significant (*P* < 0.05) differences in bacterial community composition. Next, the different samples were rescaled *via* to the “common-scale” approach (McMurdie and Holmes 2014) by means of which the proportions of all OTUs were taken, multiplied with the minimum sample size, and rounded to the nearest integer. Sampling depth of the different samples was evaluated through rarefaction curves (Figure S2) (Hurlbert 1971, Sanders 1968). The packages vegan (Oksanen et al. 2016) and phyloseq (McMurdie and Holmes 2013) were used for in-depth microbial community analysis.

A heatmap was created on the Phylum level (1% cut-off) with the *pheatmap* function (pheatmap package), and biological replicates were collated according to the method described by Connelly et al. (2017). The order-based Hill’s numbers (Hill 1973) were used to estimate differences in α-diversity between the different digesters. These Hill’s numbers represent richness (number of OTUs, H_0_), the exponential of the Shannon diversity index (H_1_) and the Inverse Simpson index (H_2_). The non-metric multidimensional scaling (NMDS) plots, based on the bacterial amplicon or methanogenic real-time PCR data, were constructed using the Bray-Curtis (Bray and Curtis 1957), Chao (Chao 1984), Jaccard, Kulczynski (Faith et al. 1987), and Mountford (Wolda 1981) distance measures. The OTUs with a significant difference (*P* < 0.05) in relative abundance between the Sulphate and Control digester were determined with the *DESeqDataSetFromMatrix* function from the DESeq2 package (Love et al. 2014)

### 2.5. Analytical techniques

Total solids (TS), total suspended solids (TSS), volatile suspended solids (VSS), volatile solids (VS), Kjeldahl nitrogen (TKN) and COD were measured according to Standard Methods (Greenberg et al. 1992). The total ammonium and sodium concentrations were measured on a 761 Compact Ion Chromatograph (Metrohm, Herisau, Switzerland), which was equipped with a Metrosep C6-250/4.0 main column, a Metrosep C4 Guard/4.0 guard column and a conductivity detector. The eluent consisted of 1.7 mM HNO_3_ and 1.7 mM dipicolinic acid. Samples were centrifuged at 3000*g* for 3 min with a Labofuge 400 Heraeus centrifuge (Thermo Fisher Scientific Inc, Merelbeke, Belgium), filtered over a 0.45 μm filter (type PA-45/25, Macherey-Nagel, Germany), and diluted with Milli-Q water to reach the desired concentration range for quantification between 1 and 100 mg L^−1^. The phosphate and sulphate concentrations were measured on a 930 Compact Ion Chromatrograph Flex Deg (Metrohm, Herisau, Switzerland), with a Metrosep A supp 5 guard, A supp 5 150/4.0 main column and a conductivity detector. Sample preparation was identical to the sodium and ammonium concentrations, as well as the concentration range for quantification. The pH and conductivity were measured with a C532 pH and C833 conductivity meter (Consort, Turnhout, Belgium), respectively. The biogas composition was measured with a Compact Gas Chromatograph (Global Analyser Solutions, Breda, The Netherlands), while the different VFA (C2-C8) were measured with a GC-2014 Gas Chromatograph (Shimadzu^®^, The Netherlands), as described in SI (S5). The total phosphorus and iron were analysed *via* Inductive Coupled Plasma Optical Emission Spectrometry – VARIAN Vista MPX, following destruction in a CEM Mars 6 Microwave Digestion System (CEM Corporation, Matthews, NC, USA).

### 2.6. Data submission

The raw fastq files that served as a basis for the bacterial community analysis were deposited in the National Center for Biotechnology Information (NCBI) database (Accession number SRP185611).

## 3. Results

### 3.1. Digester performance

An efficient start-up was obtained for all six digesters, with a steady increase in biogas production, which reached a plateau around day 25 (Figure 1), and no residual VFA (Figure 2a). From day 26 until day 42 (before sulphate addition was initiated in the Sulphate digester), a stable average methane production rate of 610 ± 60 mL CH_4_ L^−1^ d^−1^ was obtained over the six digesters. This corresponded with a COD conversion efficiency of the A-sludge to CH_4_ of 86.9 ± 8.6 %, indicating an efficient AD process.

**Figure 1.**
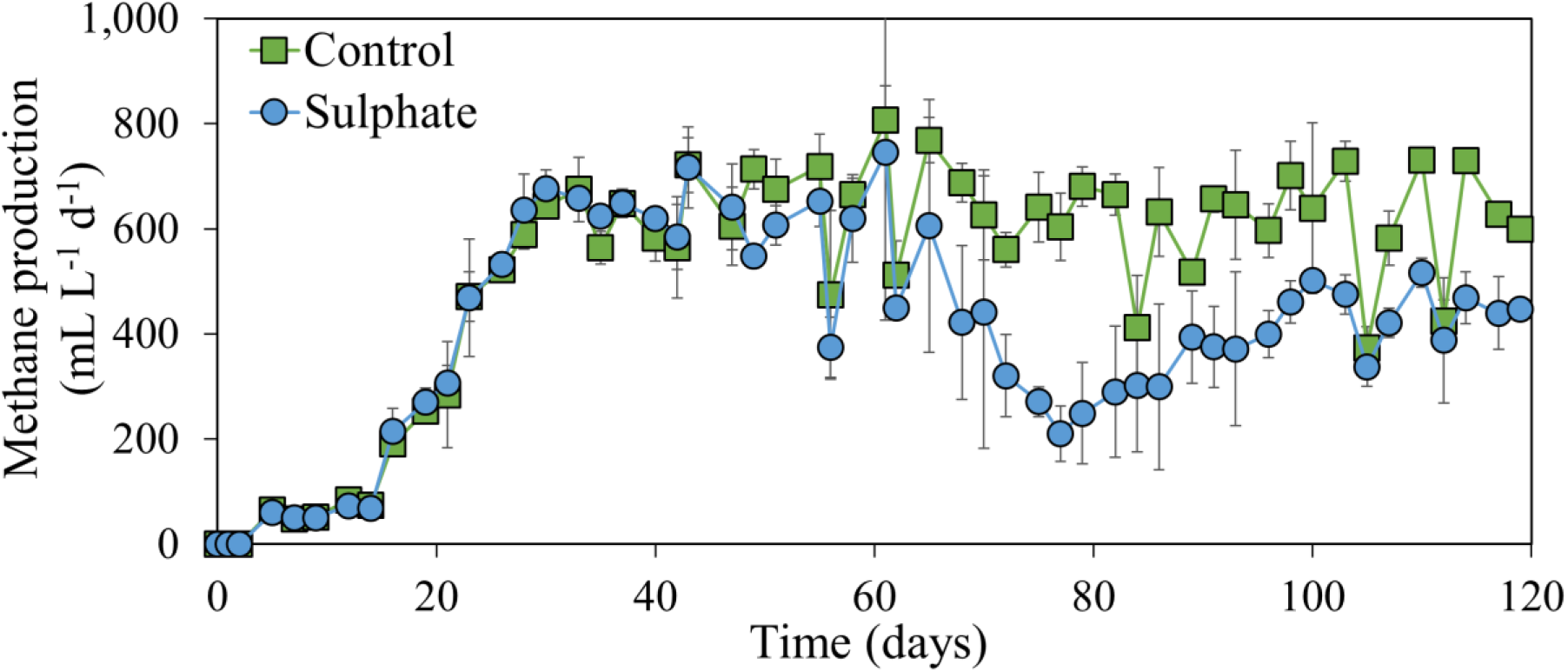
Methane production in function of time in the Control and Sulphate digester. Average values of the biological replicates (n=3) are presented, and the error bars represent standard deviations.

The Control digester continued to show stable biogas production from day 43 till day 119 (end of the experiment), with an average methane production rate of 628 ± 103 mL CH_4_ L^−1^ d^−1^, which corresponded to a COD conversion efficiency of 89.5 ± 14.6 % (Figure 1). Residual VFA concentrations did not exceed 1.5 g COD L^−1^ (Figure 2a), which corresponded to a maximum loss in COD via the effluent of 3.8 ± 1.3 % on day 91. The main VFA fractions in the Control digester from day 43 on were acetate (70.7 ± 9.5 %) and propionate (14.8 ± 11.8 %). The pH remained stable throughout the entire process, with an average value of 7.29 ± 0.10 from day 43 till day 119 (Figure 2b). Total salinity, as measured *via* conductivity, did not exceed 11.9 ± 0.1 mS cm^−1^ (Figure S3a), and the sodium concentration remained below 0.3 g Na^+^ L^−1^ (Figure S3b). The total ammonium concentration, with a maximum value of 1.34 ± 0.02 g N L^−1^ on day 77 (Figure S3c), and free ammonia concentration, with a maximum value of 32 ± 2 mg N L^−1^ on day 77 (Figure S3d), did not reach potentially inhibitory values.

**Figure 2.**
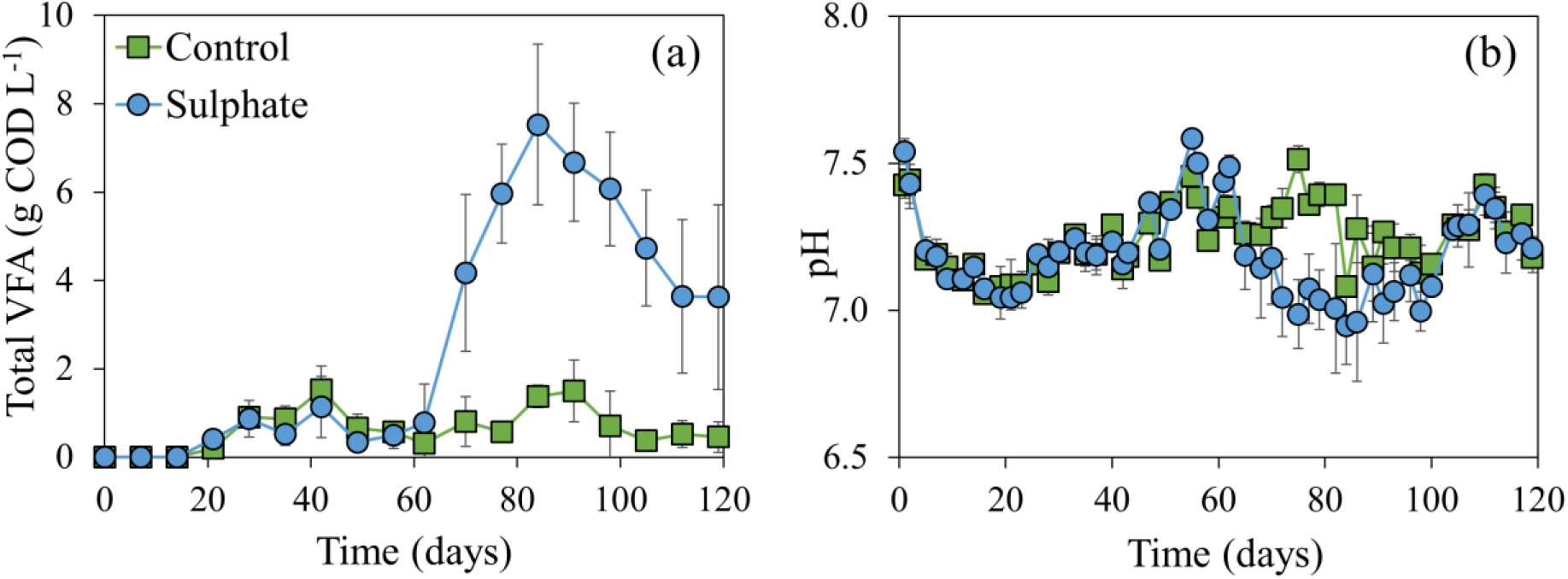
Total volatile fatty acid (VFA) concentration (a) and pH (b) in function of time in the Control and Sulphate digester. Average values of the biological replicates (n=3) are presented, and the error bars represent standard deviations.

The addition of sulphate to the feed initially did not negatively impact methane production, as an average methane production rate of 613 ± 114 mL CH_4_ L^−1^ d^−1^ and a corresponding COD conversion efficiency of 87.4 ± 16.3 % were obtained between day 43 and 61 (Figure 1). After day 61, methane production rate slowly, but steadily decreased to reach a minimum value of 210 mL CH_4_ L^−1^ d^−1^ on day 77. From day 77 on, methane production rate again slowly increased to reach a new steady state from day 98 on, with an average methane production rate of 445 ± 53 mL CH_4_ L^−1^ d^−1^ and a corresponding COD conversion efficiency of 63.5 ± 7.6 % (Figure 1). This decreasing trend in methane production and subsequent increase towards a new steady state, though at lower methane production rates, is reflected in the VFA concentration profile. The total VFA concentration increased from 0.8 ± 0.9 g COD L^−1^ on day 62 to 7.5 ± 1.4 g COD L^−1^ on day 84, but decreased again to 3.6 ± 2.1 g COD L^−1^ on day 119 (Figure 2a). This increase in VFA concentration was reflected in a decrease of the pH, though a minimum value of only 6.95 ± 0.13 was reached on day 84 (Figure 2b), which is still within the optimal range for stable AD. Total salinity was slightly higher, related to the addition of Na_2_SO_4_, with a maximum conductivity value of 15.1 ± 0.2 mS cm^−1^ on day 70 and maximum sodium concentration of 2.1 g Na^+^ L^−1^ on day 112 (Figure S3), but this was insufficient to cause direct AD process failure. In addition, neither total ammonium concentration, with a maximum value of 1.27 ± 0.01 g N L^−1^ on day 62, nor free ammonia concentration, with a maximum value of 40 ± 3 g N L^−1^ on day 62, reached potentially inhibitory concentrations.

### 3.2. Phosphate release

The key objective of this research was to evaluate to which extent sulphate reduction by sulphate reducing bacteria could assist the release of phosphate from A-sludge, related to the lower solubility product of multivalent cations with sulphides than with phosphates. In the Control digester, the phosphate concentration in the liquid phase slowly increased during the start-up from 34 ± 8 mg PO_4_^3-^ L^−1^ on day 7 to an average value of 256 ± 38 mg PO_4_^3-^ L^−1^ between day 56 and 119 (end of the experiment) (Figure 3a). Based on the total P-content of the A-sludge feedstock (Table 1), this corresponded with an average release of 12.9 ± 2.0 % of total P in the liquid phase as phosphate. As no sulphate was added to the Control digester, the residual sulphate concentration in the liquid phase remained below 25 mg SO_4_^2-^ L^−1^, except for day 0 (Figure 3b). No H_2_S could be detected in the biogas of the Control digesters, except for one replicate on day 98 (0.14% H_2_S) and day 117 (0.31 % H_2_S).

**Figure 3.**
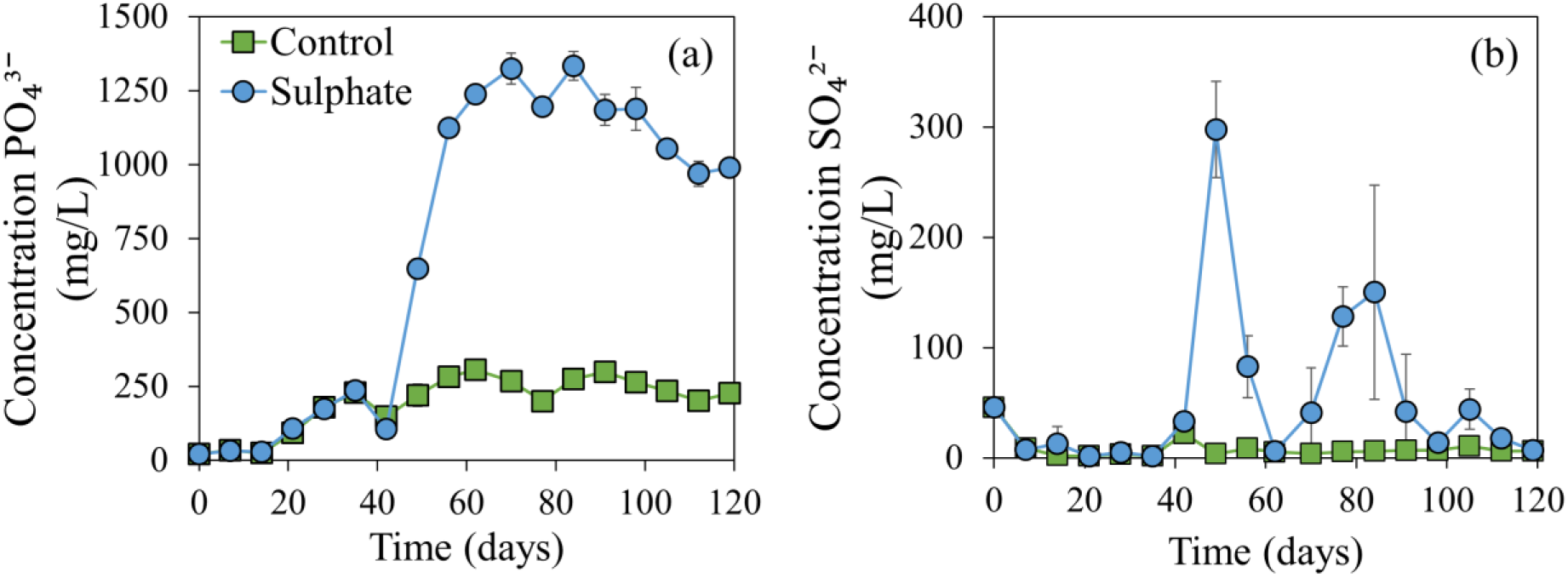
Phosphate (a) and sulphate (b) concentration in function of time in the Control and Sulphate digester. Average values of the biological replicates (n=3) are presented, and the error bars represent standard deviations.

Following the addition of Na_2_SO_4_ in the Sulphate digester on day 42, the phosphate concentration in the liquid phase increased rapidly from only 106 ± 29 mg PO_4_^3-^ L^−1^ on day 42 to 1120 ± 280 mg PO_4_^3-^ L^−1^ on day 56 (Figure 3a). An average phosphate concentration of 1160 ± 130 mg PO_4_^3-^ L^−1^ was maintained between day 56 and day 119. This corresponded with an average release of 58.7 ± 12.9 % of total P into the liquid phase as phosphate, which is a factor 4.5 higher than the Control digester. The addition of sulphate, however, also resulted in an increased residual sulphate concentration in the liquid phase. A first initial peak of 298 ± 43 mg SO_4_^2-^ L^−1^ could be observed immediately following the first sulphate addition on day 49 (Figure 3b). A second lower peak of 150 ± 97 mg SO_4_^2-^ L^−1^ was detected on day 84 after which the sulphate concentration in the liquid phase decreased to similar values as the Control digester. In contrast to the Control digester, H_2_S was detected in the biogas at multiple time points between day 89 and 119, with values up to 0.50 % H_2_S in the biogas, which corresponded with maximum 8.2 % of the sulphur added to the ingoing feedstock.

### 3.3. Microbial community analysis

#### 3.3.1. Bacterial community

Amplicon sequencing of the bacterial community yielded an average of 17,570 ± 6,505 reads and 1,431 ± 400 OTUs per sample (including singletons). Following removal of singletons and normalisation according to the common-scale approach, this was reduced to an average of 9,631 ± 183 reads and 557 ± 115 OTUs per sample. No significant differences (repeated measures ANOVA, *P* < 0.0001) in bacterial community composition were detected between the biological replicates.

The bacterial community consisted mainly of the Bacteroidetes (24.4 ± 5.6 %), Firmicutes (26.4 ± 6.5 %), Proteobacteria (13.8 ± 6.7 %) and Chloroflexi (9.1 ± 6.5 %) phyla, averaged over all samples, excluding the feedstock A-sludge (Figure 4). The A-sludge was mainly comprised of Proteobacteria (62.7 %) and Firmicutes (26.6 %) phyla. A clear increase in the Proteobacteria phylum could be observed in the Sulphate digester, reaching 17.6 ± 1.6 % on day 82 and 19.5 ± 3.1 % on day 119 relative abundance, in contrast to 7.5 ± 0.7 % on day 82 and 8.6 ± 0.9 % on day 119 relative abundance in the Control digester (Figure 4). In total 222 OTUs (0.9 % of all OTUs), considering all time points, showed a significant difference (DESeqDataSetFromMatrix, *P* < 0.05) in relative abundance between the Sulphate and Control digester. The difference in the Proteobacteria phylum between the Sulphate and Control digester mainly related to OTU00025 (*Pseudomonas, P* < 0.0001) and OTU00057 (*Rhodoferax, P* < 0.0001) (Table 3). The sulphate reducers OTU00198 (*Desulfovibrio, P* < 0.0001), OTU00219 (*Desulfobulbus, P* < 0.0001) and OTU00393 (*Desulfomicrobium, P* < 0.0001) also showed a significantly higher relative abundance in the Sulphate than in the Control digester (Table 3).

**Figure 4.**
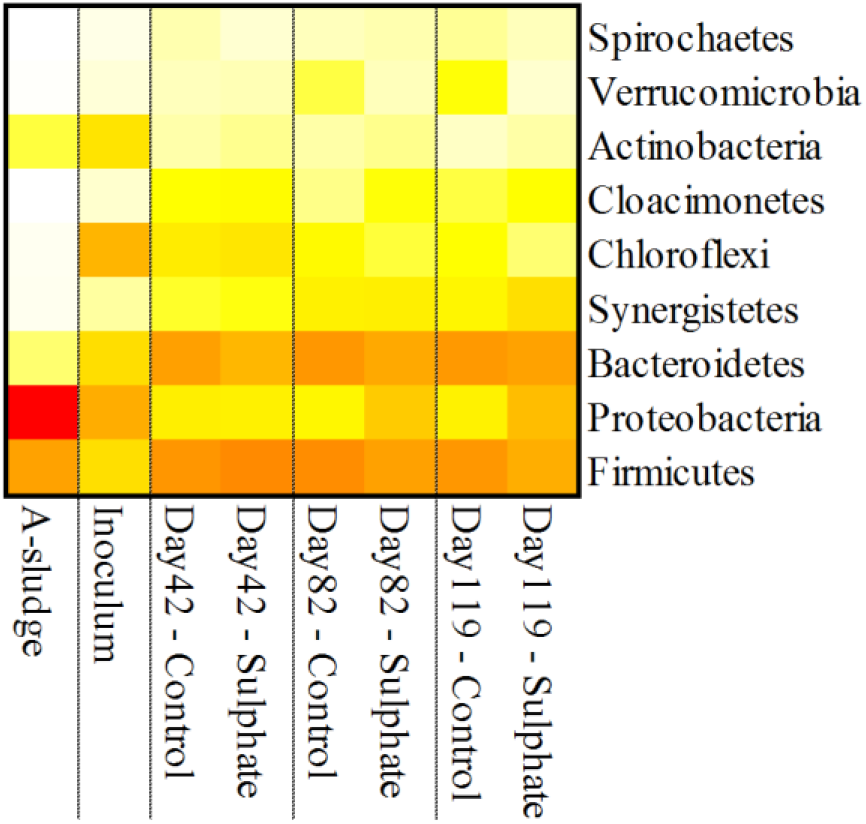
Heatmap showing the relative abundance of the bacterial community at the phylum level in the A-sludge feedstock, the inoculum and on day 42, 82 and 119 for both digesters. Weighted average values of the biological replicates are presented. The colour scale ranges from 0 (white) to 60% (red) relative abundance.

**Table 3.**
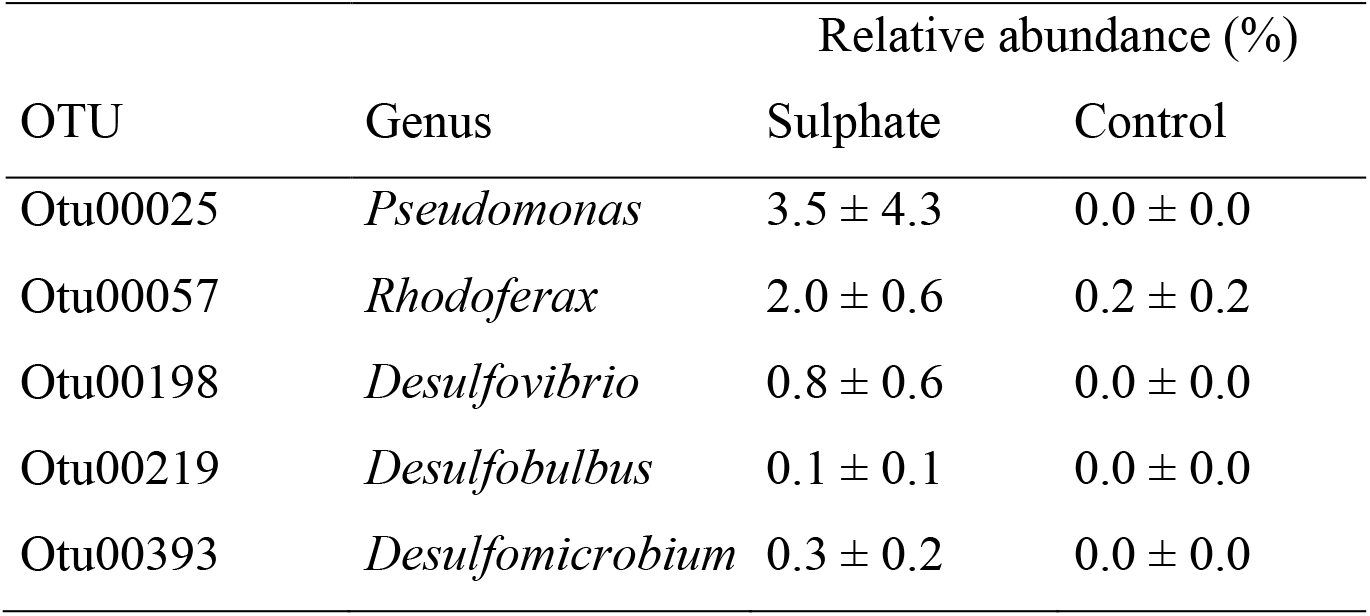
Overview of the key OTUs with their relative abundance in the bacterial community that show a significantly different relative abundance (DESeqDataSetFromMatrix, *P* < 0.0001) between the Sulphate and Control digesters.

| OTU | Genus | Relative abundance (%) |  |
| --- | --- | --- | --- |
|  |  | Sulphate | Control |
| Otu00025 | *Pseudomonas* | $3.5 \pm 4.3$ | $0.0 \pm 0.0$ |
| Otu00057 | *Rhodoferax* | $2.0 \pm 0.6$ | $0.2 \pm 0.2$ |
| Otu00198 | *Desulfovibrio* | $0.8 \pm 0.6$ | $0.0 \pm 0.0$ |
| Otu00219 | *Desulfobulbus* | $0.1 \pm 0.1$ | $0.0 \pm 0.0$ |
| Otu00393 | *Desulfomicrobium* | $0.3 \pm 0.2$ | $0.0 \pm 0.0$ |

The α-diversity analysis on the different levels of diversity (H_0_, H_1_ and H_2_) did not reveal clear differences between the digesters (Figure S4). In contrast, β-diversity analysis, based on the Bray-Curtis distance measure, revealed clear divergence in the bacterial community composition in the Sulphate digester on day 82 and 119, compared to the Control digester (Figure 5a). As sulphate addition only started on day 42, no differences in community composition were observed yet between the Control and Sulphate digester. This result was confirmed for the Jaccard, Chao, Kulczynski, and Mountford distance measures (Figure S5).

**Figure 5.**
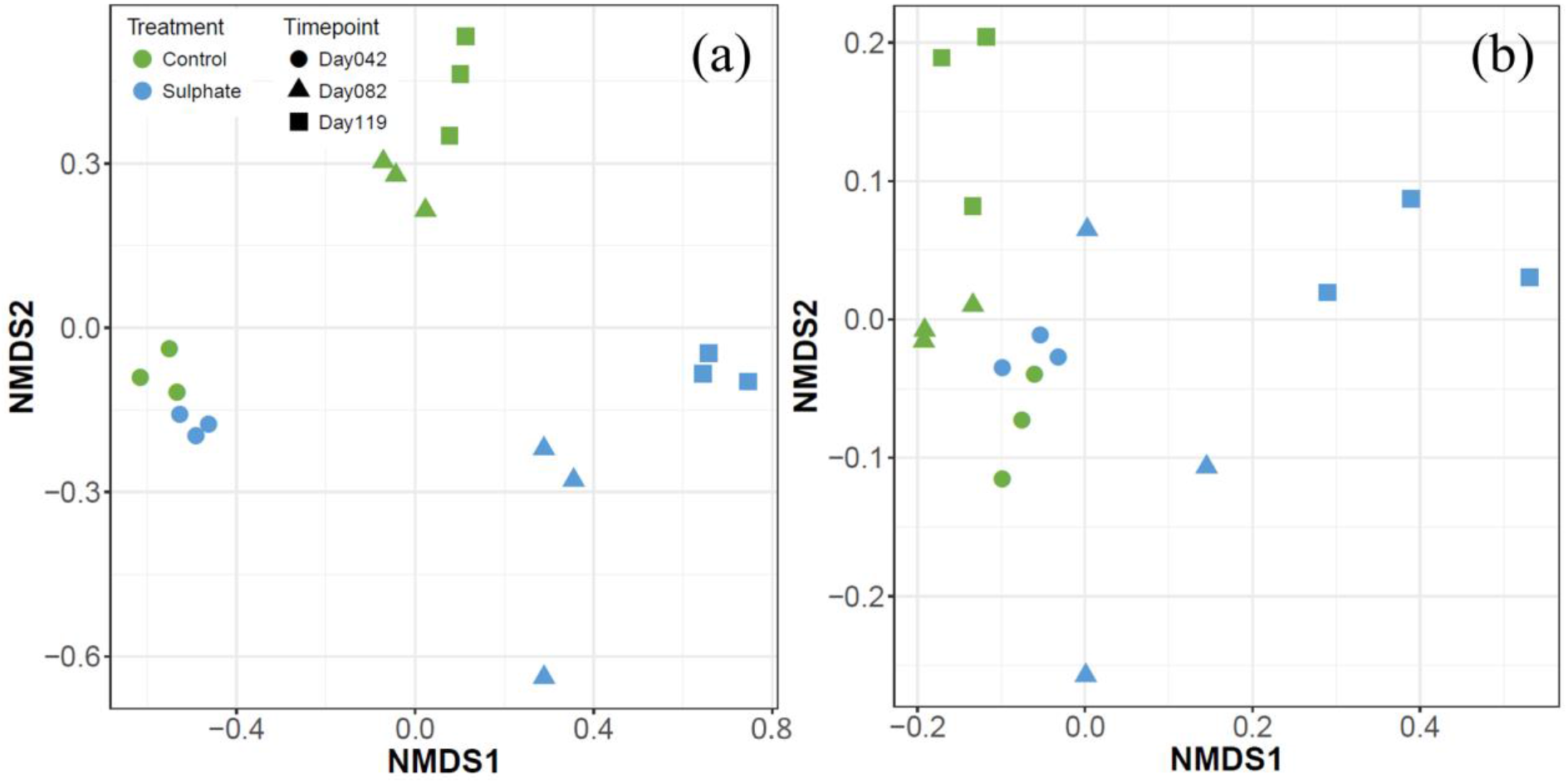
Non-metric multidimensional distance scaling (NMDS) analysis of the Bray-Curtis distance measure of the bacterial (a), based on amplicon sequencing data at OTU level (stress = 0.059), and methanogenic (b) community (stress = 0.072), based on real-time PCR data. Different colours and symbols are used for different digesters and timepoints, respectively.

#### 3.3.2. Methanogenic community

Real-time PCR analysis of the total bacteria and the different methanogenic groups revealed an overall dominance of the bacteria in absolute abundance, as the methanogens comprised only 0.26 ± 0.12 % of the microbial community, averaged over all samples, excluding the feedstock A-sludge. Both the A-sludge and Inoculum were dominated by the Methanosaetaceae (Figure 6). This was also reflected in the Control digester during the entire experiment, though an increase in relative abundance of the Methanomicrobiales could be observed at the cost of the Methanosaetaceae on day 119. The Sulphate digester showed a similar pattern as the Control digester, though on day 119, a strong increase in the relative abundance of the Methanosarcinaceae could be observed, which coincided with a reduced relative abundance of the Methanosaetaceae and Methanomicrobiales. The β-diversity analysis of the methanogenic community, based on the Bray-Curtis distance measure, confirmed this shift in the methanogenic community (Figure 5b). This shift was confirmed for the Jaccard and Kulczynski distance measures (Figure S5).

**Figure 6.**
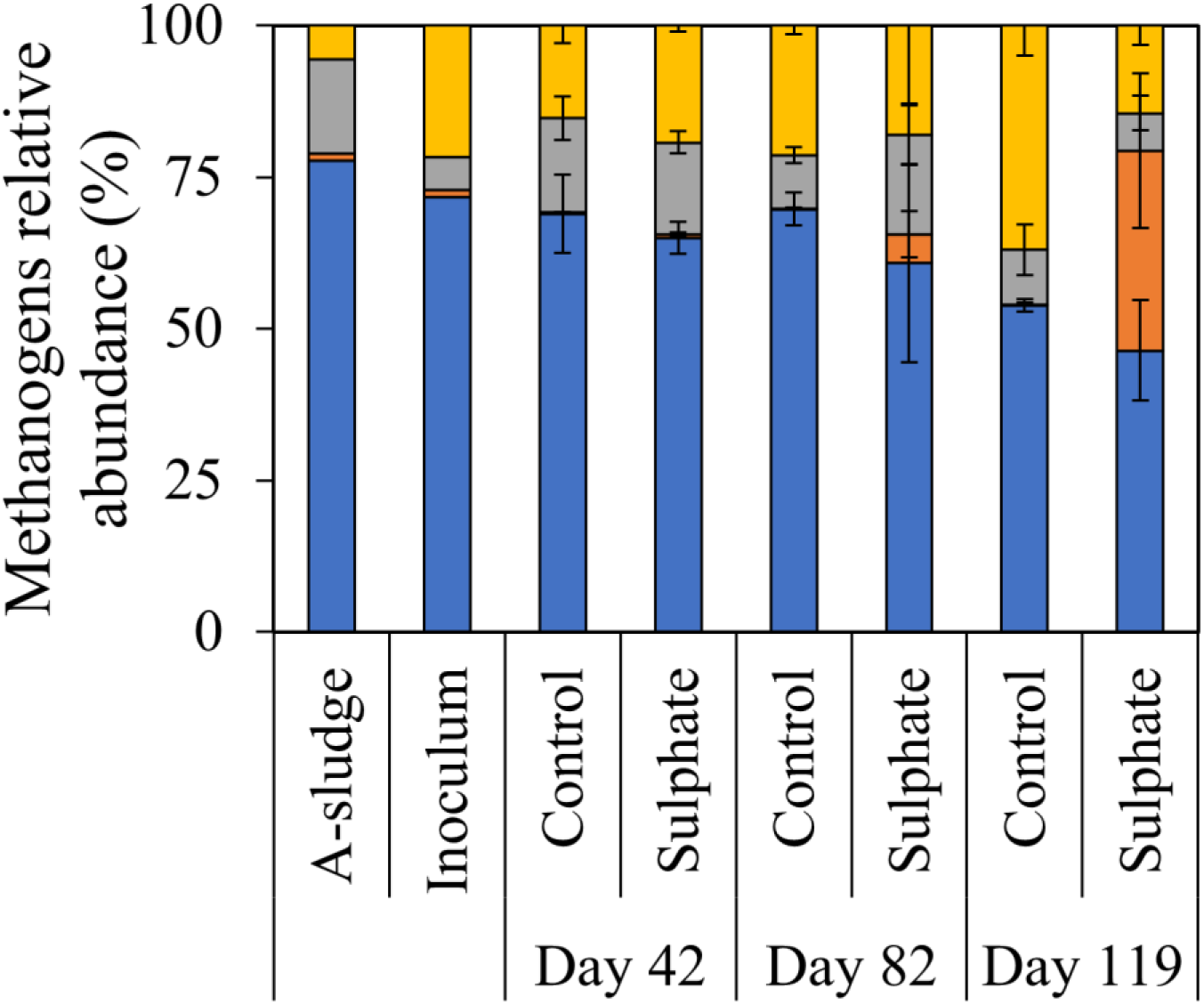
Relative abundance (%) of the Methanosaetaceae (blue, 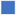), Methanosarcinaceae (orange, 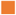), Methanobacteriales (grey, 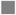) and Methanomicrobiales (yellow, 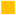) in the methanogenic community of the A-sludge feedstock, the inoculum and on day 42, 82 and 119 for both digesters. Average values of the biological replicates (n=3) are presented, and the error bars represent standard deviations.

## 4. Discussion

The stimulation of sulphate reducing bacteria by supplementing the A-sludge feedstock with sulphate enabled the release of phosphate into the liquid phase. The methanogenesis process was not strongly affected, yet, lower methane production rates and increased concentrations of residual VFA were observed. The bacterial and archaeal community demonstrated a clear shift in composition in response to the sulphate addition, with a clear increase in sulphate reducing genera.

### 4.1. Sulphate reduction enables phosphate release from high-rate activated sludge

The addition of sulphate during the AD process resulted in an increase in the phosphate concentration in the liquid phase up to a factor 4.5, compared to the Control digester. Even though, on average only 58.7 ± 12.9 % of total P in the A-sludge could be released into the liquid phase. As residual sulphate remained behind in the liquid phase and H_2_S was measured in the biogas in the Sulphate digester, this indicates that the sulphate reduction potential was not fully used, especially since an S:P molar ratio of 2 was applied. This S:P ratio of 2 was chosen to provide enough sulphides for phosphate release, yet, at the same time avoid too severe negative effects on methanogenesis, related to sulphide formation. A similar observation was made when subjecting manure to acidification either prior to or after AD, *i.e*., only about 60 % of total P could be released into the liquid phase (De Vrieze et al. 2019). Organic phosphorus, such as DNA components, cannot be released into the liquid phase by the direct effects of sulphate reduction, and requires pre-treatment methods, such as a free ammonia pre-treatment (Xu et al. 2018) or other methods that improve sludge biodegradability (Carrère et al. 2010). The effect of such a pre-treatment, however, will be limited. This is because of the already high conversion efficiency of COD, and, thus, also the organic phosphorus components, to methane in this study (86.9 ± 8.6 %), as also reported earlier (De Vrieze et al. 2013, De Vrieze et al. 2015, Ge et al. 2013, Meerburg et al. 2015). In addition, the economic feasibility of such a pre-treatment strategy depends on the increased COD conversion efficiency (Ma et al. 2011), which is anticipated to be limited, and phosphorus release. A strategy that could actively and simultaneously benefit both the COD conversion efficiency and phosphate release, such as a nitrous acid pre-treatment (Pijuan et al. 2012, Wei et al. 2018), potentially could be economically feasible.

The addition of sulphate resulted in a so-called “inhibited steady-state”, as described previously for ammonia toxicity (Nielsen and Angelidaki 2008), and salt toxicity (De Vrieze et al. 2014). Such an inhibited steady-state is characterized by elevated concentrations of residual VFA, and a lower, yet, steady methane production. This allows two different potential approaches for further process optimisation. A first option involves accurate control of sulphate dosing, based on online monitoring of residual VFA concentrations, residual sulphate concentrations, H_2_S in the gas phase, and/or methane production rates to sustain an optimal combined methanogenesis and phosphate release process. Alternatively, sulphate addition can be increased to selectively, but completely inhibit the sensitive methanogens (Karhadkar et al. 1987), thus, evolving towards fermentation with the objective to directly produce VFA instead of methane, combined with phosphate release. Both approaches require an alternative way of process engineering for targeted resource recovery of which technical and economic aspects will determine the case-specific application potential.

### 4.2. The microbial community response reflects a shift in response to sulphate addition

The accumulation of VFA and decrease in methane production in the Sulphate digester coincided with a clear shift in the microbial community. The increase in relative abundance of confirmed sulphate reducing bacteria in the Sulphate digester, relative to the Control digester is to be expected, yet, their relative abundance, except for OTU00198 in one of the biological replicates on day 119, remained below 1 % of the bacterial community. One could question the involvement of these sulphate reducing bacteria in the overall process, yet, given their ability to reduce sulphate and complete absence in the different biological replicates of the Control digester on day 82 and 119, their involvement in the sulphate reduction process is apparent. The potentially important role of low-abundant OTUs in AD has been indicated frequently (Guo et al. 2015, Theuerl et al. 2018, Vanwonterghem et al. 2016), and is also reflected in the present study, with an overall low relative abundance of the methanogens, which are nonetheless essential in the AD process.

The methanogenic community in the Sulphate digester showed a clear shift towards an increased relative and absolute abundance of the Methanosarcinaceae at the expense of the Methanosaetaceae, in contrast to the Control digester. The overall higher tolerance of *Methanosarcina* sp., compared to *Methanosaeta* sp., to different stressors in AD (Conklin et al. 2006, De Vrieze et al. 2012), explains this shift, in response to the formation of sulphides. A similar shift has been observed in other studies, in response to multiple stressors, although the methane production pathway by the *Methanosarcina* sp., *i.e*., either acetoclastic or hydrogenotrophic methanogenesis, may vary (De Vrieze et al. 2012, Lins et al. 2014, Lu et al. 2013, McMahon et al. 2001, Poirier et al. 2016, Venkiteshwaran et al. 2016). The apparent “inhibited steady state” in this study, as also observed previously, thus, seems to be a consequence of the shift from a *Methanosaeta* sp. to *Methanosarcina* sp. domination, and reflects their different metabolic features.

### 4.3. The cost of phosphate release through sulphate reduction

The release of phosphate into the liquid phase due to sulphide formation was up to a factor 4.5 higher than when no sulphate was added to the feedstock. This enables an integrated valorisation of the A-sludge, with combined energy (through biogas) and nutrient (through phosphate) recovery. For this purpose, the “inhibited steady-state” phase (day 89-119) of the Sulphate digester (428 ± 53 mL CH_4_ L^−1^ d^−1^) was compared to the Control digester (611 ± 110 mL CH_4_ L^−1^ d^−1^) during that same period, which corresponded with a 29.9 ± 15.3 % lower methane production in the Sulphate compared to the Control digester. An electrical efficiency of 40 % for the CHP unit (Deublein and Steinhauser 2008, Szarka et al. 2013), a higher heating value of 9.95 kWh m^−3^ CH_4_, a 95 % recovery of methane from the digester, and a hydraulic retention time of 20 days were assumed. At a current electricity market price of € 0.10 kWh^−1^ (De Vrieze et al. 2016), this amounts € 126 m^−3^ year^−1^ for the Control and € 89 m^−3^ year^−1^ for the Sulphate digester per unit digester volume for electricity from biogas, thus, a deficit of € 37 m^-3^ year^−1^ related to sulphate reduction. Projected electricity market prices of € 0.03 kWh^−1^ by 2020-2025, and even down to € 0.01 kWh^−1^ by 2030-2040 (Fraunhofer 2015, van Wijk et al. 2017) reduce this deficit to € 8 & 3 m^−3^ year^−1^, respectively. Based on the market price of phosphate rock of € 350-1200 tonne^−1^ P since 2010 (Mayer et al. 2016), this yields a potential revenue of € 0.5 m^−3^ year^−1^ for the Control and 2.3 m^−3^ year^−1^ for the Sulphate digester at a value of € 350 tonne^−1^ P. At a value of € 1200 tonne^−1^ P, this revenue increases to € 1.8 & 7.9 m^−3^ year^−1^ for the Control and Sulphate digester, respectively. This indicates that the deficit due to the decrease in methane production can be at least partially covered by the revenue from phosphorus recovery, but additional resource recovery strategies and/or advanced control of the sulphate reduction process will be essential.

The integrated approach of this study that makes use of sulphate reduction during AD for combined energy and phosphorus recovery faces additional challenges. First, the release of phosphate into the liquid phase allows subsequent recovery, yet, phosphate recovery technologies, such as struvite precipitation, result in additional costs and product quality issues. For example, phosphorus recovery from pig manure through struvite crystallisation coincides with an electricity cost of € 0.5 tonne^−1^ pig manure (De Vrieze et al. 2019, Flotats et al. 2011). Second, the increase in H_2_S in the biogas requires additional desulfurization, for which both physicochemical and biological techniques can be used (Abatzoglou and Boivin 2009), to a recommended value < 0.025 % (Weiland 2010) before it can be sent to the CHP unit. Third, the presence of residual VFA in the liquid phase, with an average value of 4.9 ± 1.4 g COD L^−1^ between day 89-119, necessitates additional (aerobic) effluent polishing.

These challenges can also be considered opportunities. The H_2_S in the biogas can be recovered as elemental sulphur *via* different techniques (Pandey and Malhotra 1999). The phosphate and residual VFA can be recovered in a combined electrodialysis or membrane electrolysis approach (Andersen et al. 2014, De Vrieze et al. 2018). Alternatively, the AD process can be shifted from methanogenesis to direct VFA production, thus, avoiding biogas desulfurization and focusing on the liquid phase. Hence, an integrated approach that combines the release and recovery of phosphorus with other resource recovery strategies for the valorisation of organic waste and side streams could find its way towards future full-scale applications.

## 5. Conclusions

The exploitation of sulphate reduction for the release of phosphorus from energy-rich sludge increased the phosphate concentration in the liquid phase with a factor 4.5. The sulphate reduction process pushed the anaerobic digestion process to an inhibited steady state, as reflected in both operational and microbial parameters, with reduced, yet, stable methane production rates. Phosphate can be recovered in an economically feasible way, but only in combination with energy, organics or other resources for integrated valorisation of energy-rich sludge.

## Acknowledgments

Jo De Vrieze is supported as postdoctoral fellow by the Research Foundation Flanders (FWO-Vlaanderen). The authors would like to thank Tim Lacoere for his assistance with the molecular analysis, and Cindy Law, Amanda Luther and Inka Vanwonterghem for carefully reading the manuscript. The authors also kindly acknowledge Harry Vrins from Waterschap Brabantse Delta for his assistance with the A-sludge collection.

